# Enantioselective Protein Affinity Selection Mass Spectrometry (E-ASMS)

**DOI:** 10.1101/2025.01.17.633682

**Authors:** Xiaoyun Wang, Jianxian Sun, Shabbir Ahmad, Diwen Yang, Fengling Li, U Hang Chan, Hong Zeng, Conrad V. Simoben, Stuart R. Green, Madhushika Silva, Scott Houliston, Aiping Dong, Albina Bolotokova, Elisa Gibson, Maria Kutera, Pegah Ghiabi, Ivan Kondratov, Tetiana Matviyuk, Alexander Chuprina, Danai Mavridi, Christopher Lenz, Andreas C. Joerger, Benjamin D. Brown, Richard B. Heath, Wyatt W. Yue, Lucy K. Robbie, Tyler S. Beyett, Susanne Müller, Stefan Knapp, Rachel Harding, Matthieu Schapira, Peter J. Brown, Vijayaratnam Santhakumar, Suzanne Ackloo, Cheryl H. Arrowsmith, Aled M. Edwards, Hui Peng, Levon Halabelian

## Abstract

We report an enantioselective protein affinity selection mass spectrometry screening approach (E-ASMS) that enables the detection of weak binders, informs on selectivity, and generates orthogonal confirmation of binding. After method development with control proteins, we screened 31 human proteins against a designed library of 8,210 chiral compounds. 16 binders to 12 targets, including many proteins predicted to be “challenging to ligand”, were discovered and confirmed in orthogonal biophysical assays. 7 binders to 6 targets bound in an enantioselective manner, with *K*_D_ values ranging from 3 to 20 µM. Binders for four targets (DDB1, WDR91, WDR55, and HAT1) were selected for in-depth characterization using X-ray crystallography. In all four cases, the mechanism for enantioselectivity was readily explained. We conclude E-ASMS can be used to identify and characterize selective and weakly-binding ligands for novel protein targets with unprecedented throughput and sensitivity.

## Main

Bioactive small molecules are invaluable reagents in basic research and applied fields such as biomedicine, agriculture and microbiology. The discovery of bioactive small molecules typically begins with screening of synthetic chemical or natural product libraries. Often, libraries of synthetic chemical or natural product libraries are screened in high-throughput or phenotypic screening^1–3^ to identify compounds that alter biochemical or cellular functions. Alternatively, target-centric approaches can be employed, such as methods that measure binding to the target protein directly *in vitro*, including fragment-based lead discovery (FBLD)^4–6^, DNA-encoded library (DEL) selection, or affinity selection mass spectrometry (AS-MS)^7^. A significant challenge across all screening strategies is to develop orthogonal assays to distinguish true hits from false positives. Indeed, particularly when the initial hits bind weakly (> 10 µM *K*_D_), this hit verification process often demands more time and resources than the initial screen. As a result, it has proven impractical for the discovery of chemical ligands for large numbers of proteins.

Small molecules with a stereocenter may bind to their targets in an enantioselective way^10^, which can be leveraged to identify potential stereoselective protein binders when screening libraries containing chiral compounds, as well as to develop negative controls for chemoproteomics studies and for experiments with chemical probes^11–13^. To exploit this phenomenon, we developed a scalable orthogonal screening strategy, termed “Enantioselective Protein Affinity Selection Mass Spectrometry (E-ASMS)”, to both identify and characterize chemical ligands for previously unliganded proteins.

## Results

### Design and benchmark an E-ASMS platform

#### Method concept

The E-ASMS screening concept workflow (Fig. 1a) is a variation of previous methods used for bioactive natural product discovery^14–16^, by including the orthogonal enantioselectivity evidence. First, pools of ~600 compounds from an 8,210 member screening library, comprising racemic mixtures of drug-like compounds at a concentration of 0.1 µM, are incubated with purified and quality-controlled polyhistidine-tagged proteins immobilized on magnetic nickel beads. At this concentration, all compounds are far below their solubility limit in aqueous solution. To embed a measure of selectivity in the screen, 8-12 different proteins are screened in parallel at a concentration of 1 µM in solution. After washing the beads, compounds that remain bound to the proteins and the beads, which comprise a combination of real and nonspecific binders, are eluted in methanol. A small aliquot of each methanol eluate is subject to liquid chromatography (LC) followed by high-resolution mass spectrometry (MS) to identify any compounds in the eluates, and to estimate their abundance relative to the eluates from other targets screened in parallel. Our concept is that some measure of binding specificity can be inferred by observing strong binding of a compound to one protein but not to the other proteins screened in parallel. For the compounds that appear to bind one target preferentially over the others, another aliquot from the same methanol eluate is analyzed by chiral chromatography, and the ratio of the two enantiomers in the protein eluate is assessed. If one of the two enantiomers is enriched in the protein eluate compared to their ratio in the screening library, then this provides orthogonal evidence for compound binding.

**Fig. 1.**
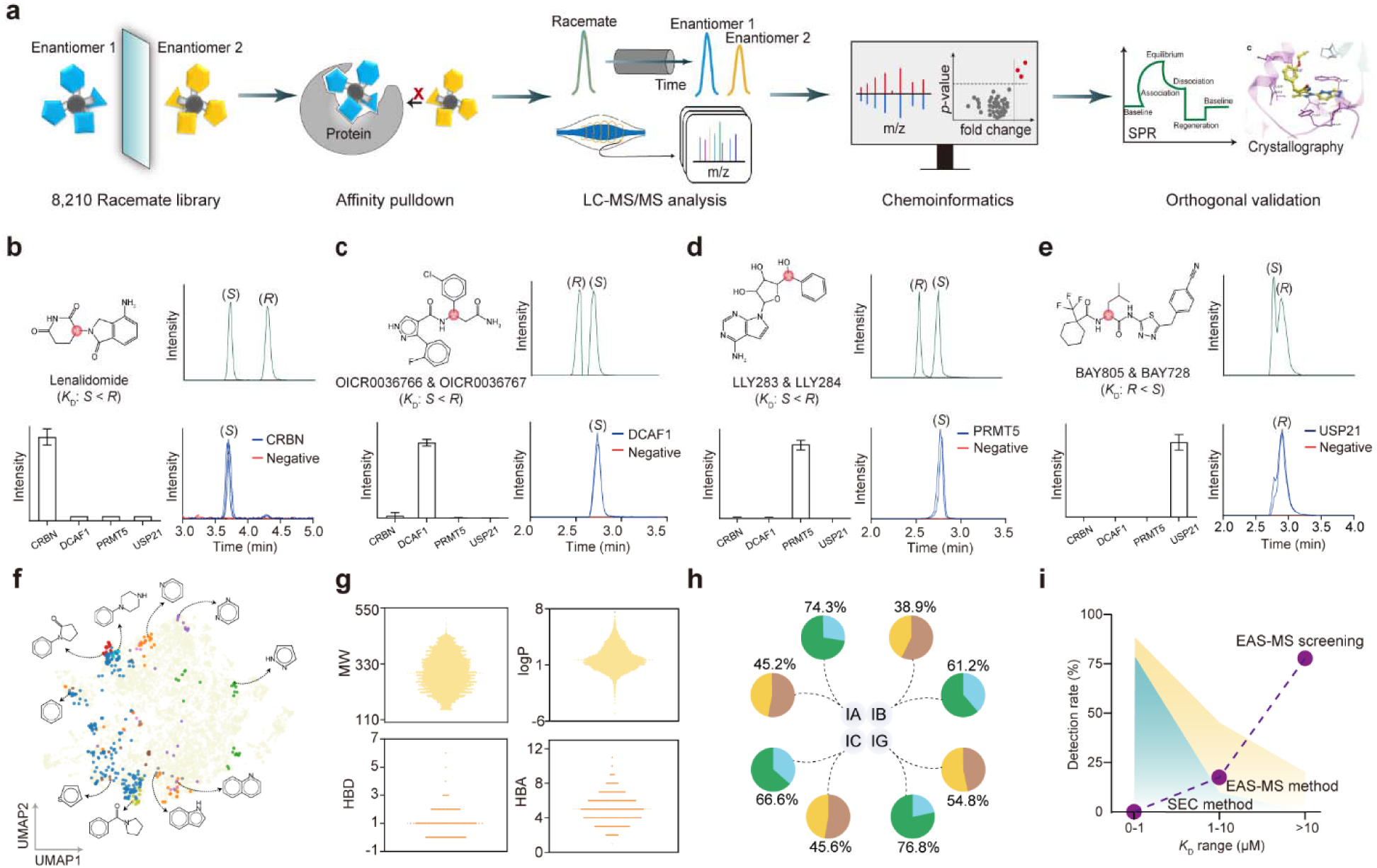
Development of the E-ASMS platform for scalable protein ligandability discovery. (**a**) Schematic representation of the E-ASMS screening workflow. (**b-e**) Benchmarking the E-ASMS platform using four positive control ligands binding to CRBN, DCAF1, PRMT5, and USP21, respectively. (**f**) Chemical space of the E-ASMS library. A two-dimensional uniform manifold approximation and projection (UMAP) visualization of SMILES descriptor. The displayed structures represent the top 10 scaffolds within the E-ASMS chemical library. (**g**) Molecular properties of the E-ASMS chemical library: molecular weight (MW), logP, hydrogen bond acceptors (HBA) and donors (HBD). **(h)** Chiral separation of the E-ASMS library using four chiral columns (IA, IB, IC, and IG), and two mobile phases (acetonitrile/methanol, yellow; water/methanol, green). **(i)** Detection rate of 109 ligands against 8 proteins by SEC and E-ASMS methods, respectively. Purple dots represent the *K*_D_ distribution of chemical ligands discovered in the present E-ASMS screening against 31 proteins.

#### Proof of concept for E-ASMS with known enantioselective protein-ligand pairs

To pilot the E-ASMS concept, we selected four chiral compounds known to bind to their respective protein partners in an enantioselective manner. The first test case was lenalidomide, a chiral analogue of thalidomide that binds enantioselectively to the cereblon (CRBN) protein^17–19^. As shown in Fig. 1b, when screened against four proteins (CRBN, DCAF1, PRMT5, and USP21), lenalidomide was selectively enriched in the methanol eluate from CRBN beads; no significant enrichment was observed on beads containing any of the three other human proteins, which served as negative controls. The CRBN methanol eluate was then subjected to chiral chromatography. We recovered (*S*)-lenalidomide in the eluate from CRBN-containing beads at far greater quantities compared to its (*R-*) enantiomer, despite both being present in equal amounts in the input. Enantioselective enrichment was also observed for (*S*)-OICR0036766, (*S*)-LLY283, and (*R*)-BAY805 to their cognate protein targets: DCAF1^20^, PRMT5^21^, and USP21^22^ (Fig. 1c, 1d, and 1e). *Racemate library design and characterization*: With the proof of concept in hand, we set out to apply the method to novel proteins in screening mode. To this end, we assembled a chemically diverse screening library with drug-like properties and high MS sensitivity, comprising 8,210 commercially available racemates (Supplementary Fig. 1 and Supplementary Table 1). The E-ASMS library was designed to maximize chemical diversity, encompassing 7,307 distinct scaffolds (Supplementary Fig. 2), with the top 10 scaffolds highlighted in Fig. 1f. The designed E-ASMS library adheres to Lipinski’s rule of 5 (*e.g.,* MW, 119 – 478 Da; logP, 1.65 ± 1.42; nHBD, 0 – 6; nHBA, 1 – 11; Figure 1g). Chromatographic method(s) was developed to resolve the enantiomers for as many of the racemates as was practical. To accomplish this, each of the 8,210 racemic mixtures was subject to chiral liquid chromatography under 8 different conditions (4 columns × 2 LC methods). In total, ~7,000 enantiomer pairs could be resolved under one of these conditions. The most effective chromatographic steps involved using IG and IA columns (Fig. 1h). Thus, for any of these ~7,000 compounds, it is possible to compare the ratio of enantiomers in the screening library and the protein eluant, as described in the workflow outlined above (Fig. 1a).

#### E-ASMS identifies low affinity binders

Rapid size-exclusion chromatography (SEC) is the most common approach used to resolve protein binders and non-binders in AS-MS library screening^7^. However, this method is known to be limited by the inability to detect weakly bound compounds, perhaps due to the challenges in retaining compounds with fast off-rates during chromatography. We explored whether the E-ASMS platform in which the protein is concentrated on beads and employs rapid wash steps, had the potential to enhance the sensitivity of AS-MS screens. Using 109 previously characterized ligands across 8 proteins (Supplementary Table 2), we tested the sensitivity of E-ASMS. The E-ASMS platform successfully captured nearly all high-affinity (*K*_D_ < 1 µM) ligands, half of the moderate-affinity (1 µM ≤ *K*_D_ < 10 µM) ligands, and 20% of the low-affinity (*K*_D_ ≥ 10 µM) ligands (Fig. 1i) without substantially increasing the false positive rate. In contrast, all moderate- and low-affinity ligands were lost by the SEC-coupled AS-MS approach. The apparent sensitivity of E-ASMS is a significant advantage for chemical ligand discovery, or to assess the ligandability of a new protein, as binders can be rapidly identified using a relatively small chemical library and minimal amounts of protein in a protein-agnostic manner.

### Application of the E-ASMS platform to novel targets

With evidence of increased sensitivity and an ability to potentially provide orthogonal evidence for specificity using enantioselectivity, we set out to explore the application of the E-ASMS platform to new proteins. We selected 31 human proteins with diverse biological functions, 3D structures, and with varying levels of known or predicted ‘ligandability’ (Fig. 2a) as assessed by their drug-like density (DLID) scores^23^. We purposely included several targets (*e.g.,* WDR91 and DCAF1) known to be ‘ligandable’ ^20,24,25^. Among the 31 proteins, 3 were predicted to have high ligandability (DLID > 1), 11 of medium ligandability (0.5 < DLID < 1), and 16 of low ligandability (DLID < 0.5) (Supplementary Fig. 3).

**Fig. 2.**
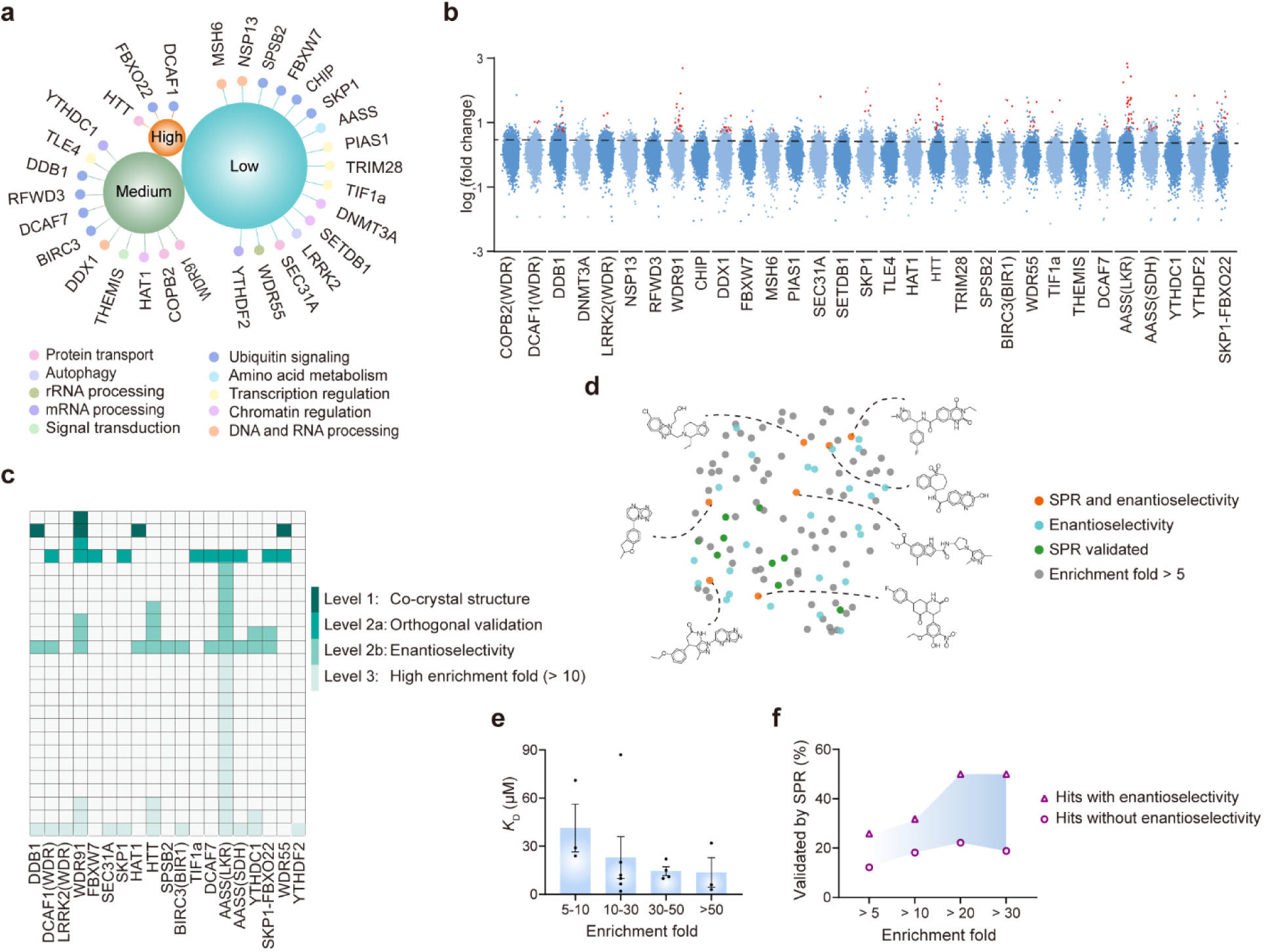
Scalable chemical ligand profiling across 31 diverse proteins. (**a**) Overview of the 31 screened proteins. ‘High’: drug-like density (DLID) > 1, ‘Medium’: 0.5 < DLID < 1, ‘Low’: DLID < 0.5. (**b**) Scatter plot of all E-ASMS library compounds interacting with 31 proteins. Dots above the line represent hits with enrichment fold > 5. Red dots indicate hits with enrichment fold > 5 and *p*-value < 0.05. (**c**) Confidence levels of all identified hits according to orthogonal evidence. (**d**) UMAP visualization of the chemical space for hits with enrichment fold > 5. (**e**) Correlation between *K*_D_ and enrichment fold. (**f**) SPR validation rate of hits with or without enantioselectivity, across different enrichment fold ranges.

The 31 proteins were grouped in 4 batches of 8 proteins and screened against 14 pools of ~600 compounds. In total, 118 compounds (candidate binders) exhibited > 5-fold enrichment for one protein compared to the others screened in parallel, corresponding to an average hit rate of 0.05% (Fig. 2b and Supplementary Table 3). Candidate binders were detected for all three high ligandable targets, seven of the eleven targets with medium ligandability scores, and nine of the sixteen targets predicted to be the most challenging. Interestingly, these binders were distributed across the entire chemical space of the E-ASMS library (Supplementary Fig. 4).

To confirm binding, we tested the binding of 101 candidate binders to each of the 19 targets using orthogonal biophysical methods. We used surface plasmon resonance (SPR), a technique that requires relatively small amounts of protein and can provide quantitative orthogonal confirmation of binding in a pocket-agnostic manner. The binding of 16 hits to 12 targets was confirmed by SPR, ranging from 1-4 hits per target and with *K*_D_ values ranging from 2 to 87 µM (Figure 2c, 2d, and Supplementary Table 3). Notably, 13 of the 16 SPR-confirmed hits exhibited low-affinity (*K*_D_ > 10 µM) (Fig. 1i), which might have been overlooked by the conventional SEC-coupled AS-MS method. According to orthogonal evidence, we categorized 118 candidate binders into four confidence levels (Fig. 2c). Interestingly, the degree of enrichment in the primary E-ASMS screen showed a strong correlation with the *K*_D_ values determined for all SPR-validated hits (Fig. 2e), demonstrating the potential application of the E-ASMS method for semi-quantitative read-outs.

Ideally, SPR assay methods are developed using a positive control binder, which we lacked for nearly all of the proteins. Accordingly, positive results by SPR can be interpreted, but negative results are inconclusive. The 85 candidate binders that were unable to be confirmed using SPR are not necessarily false positives for three reasons. First, they might not be able to be detected because the assay for the protein target could not be optimized using a positive control compound. Second, they might also be *bona fide* binders with affinities beyond the sensitivity of SPR under the non-optimized solution conditions used. Third, the compounds might be insoluble at the concentrations required and conditions used for SPR (up to 0.2 mM of the compound) and might confound the read-outs^26^. The third possibility is a common issue when attempting to characterize weakly binding compounds by any biophysical method.

### Enantioselective enrichment provides orthogonal evidence of compound binding

Our E-ASMS method, which creates an effective protein concentration of ~100 µM on the beads, enables the screening of compounds at a concentration of 100 nM, far below their solubility limits. In support of solubility being a confounding issue for SPR orthogonal validation, we observed that SPR confirmation rates improved significantly for E-ASMS hits with higher fold enrichment values. For E-ASMS hits without enantioselectivity that were enriched > 5 fold, the SPR-confirmation rate was 12.2%; this increased to 22.2% for more potent hits with fold enrichment values > 20 (Fig. 2f). For the more potent binders that were enantioselective enrichment, the SPR confirmation rate increased further to 50%. Even subject to the caveats of SPR, this trend clearly demonstrates the potential value of enantioselective enrichment in hit confirmation.

Among the 118 candidate binders identified in the screen of 31 proteins, clear enantioselective binding was detected for 32 binders to 14 targets (Fig. 2c). Of these targets, we were successful in developing an SPR assay for 6, and could calculate *K*_D_ values for 7 binders (Fig. 2d), with affinities ranging from 3-20 µM (Supplementary Figs. 5-9). There were 8 targets and 25 candidate binders for which we were unable to generate convincing binding data using either SPR under standard conditions or with other biophysical methods, such as differential scanning fluorimetry or ^19^F-NMR. At this point, we cannot be certain if these 25 enantioselective candidate binders are false positives or are true positives that are challenging to assay.

### In-depth characterization of enantioselective binding to four targets

Four targets – DDB1, WDR91, WDR55, and HAT1 – predicted to have ‘medium’ or ‘low’ ligandability scores, as measured by their DLID scores (Fig. 2a and Supplementary Fig. 3), were selected for further analysis.

*Damage-specific DNA-binding protein 1 (DDB1)* is a multidomain protein involved in protein homeostasis^27^. We screened histidine-tagged DDB1 and other proteins in parallel against the racemate library and identified a compound (XS381952) specifically enriched in the methanol eluate from DDB1 beads (Fig. 3a). When XS381952 in the DDB1-bead eluate was analyzed by chiral chromatography, one of the two enantiomers was clearly enriched compared with the ratio of the enantiomers in the starting library, providing compelling evidence of specific binding (Fig. 3b). To rigorously characterize the binding, we resolved and characterized the two enantiomers of XS381952 using preparative chiral chromatography and electronic circular dichroism (ECD) spectroscopy (Supplementary Fig. 10), and measured their binding to DDB1 using SPR. We found that (*S*)-XS381952 bound DDB1 with a *K*_D_ of 2 µM (Supplementary Fig. 5), significantly more potent than its (*R*)-counterpart (estimated *K*_D_ > 73 µM). To elucidate the mechanism of binding, we determined the crystal structure of (*S*)-XS381952 with DDB1 (Supplementary Fig. 11a and Supplementary Table 4), and found that (*S*)-XS381952 is sandwiched between two aromatic residues, W1047 and Y1114, via π-stacking interactions (Fig. 3c). The enantioselectivity was explained by the positioning of the ethoxyphenyl moiety linked to the chiral carbon, which adopted a nearly 90° angle from the compound backbone, orienting toward V1132 and stabilizing against the 1129-1140 α-helix of DDB1. The (*R*)-enantiomer would not be able to fit into the binding site due to steric hindrance.

**Fig. 3.**
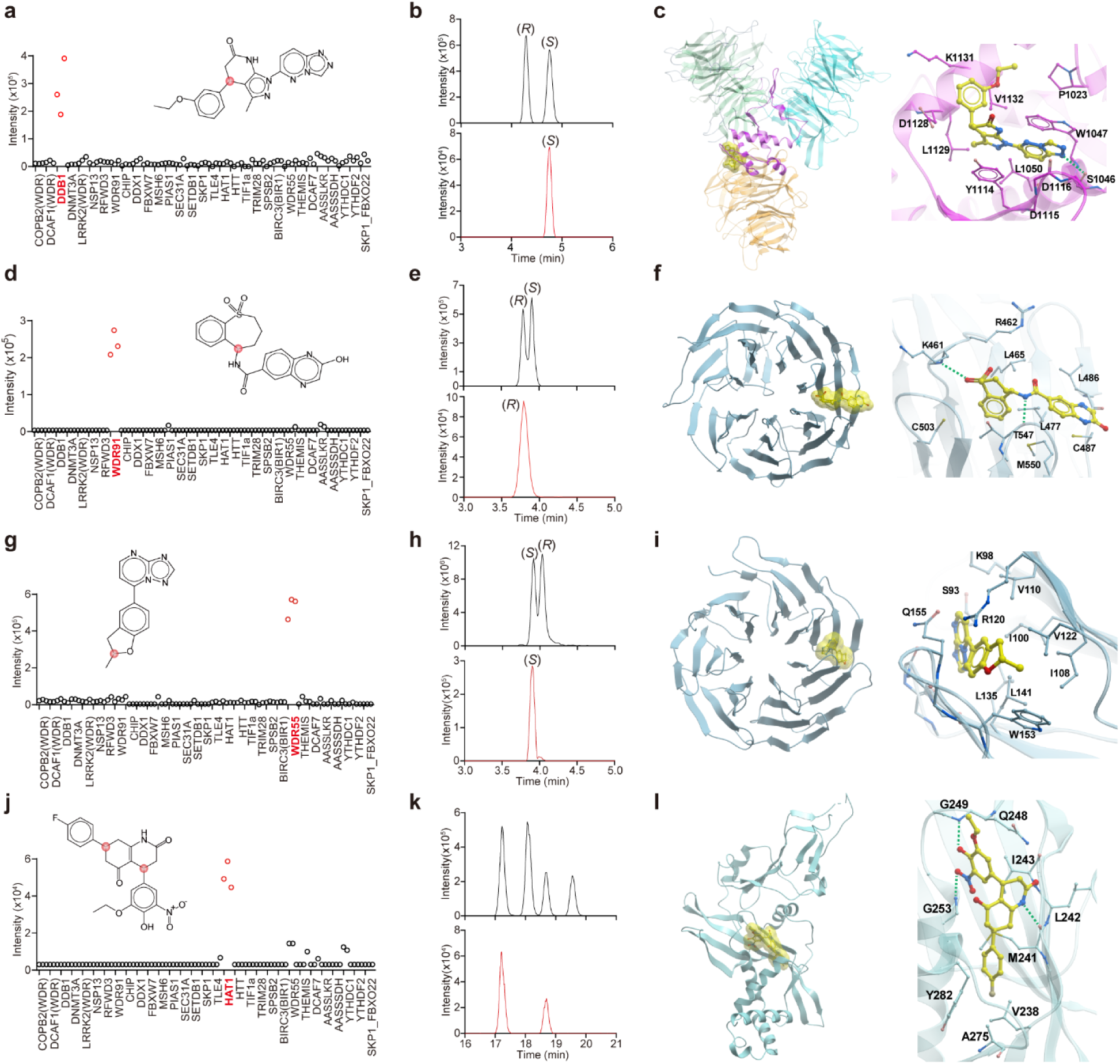
Identification of enantioselective hits for DDB1, WDR91, WDR55, and HAT1. (**a**) Scatter plot showing XS381952 pulled down by DDB1. (**b**) Chiral chromatograms of the XS381952 enantiomers before (upper) and after E-ASMS (bottom). (**c**) Co-crystal structure of DDB1 with XS381952. Hydrogen bond between the protein and the ligand is shown as green dashed line. (**d**) Scatter plot showing XS838489 pulled down by WDR91. (**e**) Chiral chromatograms of the XS838489 enantiomers. (**f**) Co-crystal structure of WDR91 with XS838489. Hydrogen bonds between the protein and the ligand are shown as green dashed lines. (**g**) Scatter plot showing XS381774 pulled down by WDR55. (**h**) Chiral chromatograms of the XS381774 enantiomers. (**i**) Co-crystal structure of WDR55 with XS381774. (**j**) Scatter plot showing XS380871 pulled down by HAT1. (**k**) Chiral chromatograms of the XS380871 enantiomers. (**l**) Co-crystal structure of HAT1 with XS380871. Hydrogen bonds between the protein and the ligand are shown as green dashed lines.

*WD40 repeat containing protein 91 (WDR91)* is a 747-residue protein with a C-terminal WDR domain that plays a critical role in endosomal maturation^29^. After screening the WDR domain of WDR91 with the racemate library, four compounds were enriched in WDR91-bead eluates compared with the other proteins screened in parallel (Supplementary Fig. 6). Two of these hits, XS838489 (Fig. 3d and 3e) and XS837729, bound in an enantioselective manner. The remaining two hits, XS381295 and XS381186, did not show enantioselectivity in its binding. The binding of all four hits was characterized by SPR, and three hits displayed single digit micromolar *K*_D_ values (Supplementary Fig. 6), while the fourth hit, XS381186, displayed 29 µM *K*_D_ (Supplementary Fig. 6). To elucidate the binding mode of racemic XS838489, we co-crystallized it with the WDR domain of WDR91 (residues 392-747), revealing the (*R*)-enantiomer bound to WDR91 (Fig. 3f, Supplementary Fig. 11b, and Supplementary Table 4). This binding pocket lies between two β-propellers and is surrounded by hydrophobic residues, including L465, L467, L477, and A459. The amide nitrogen linked to the chiral carbon forms a hydrogen bond with the backbone oxygen of T547, favoring the (*R*)-enantiomer for binding. The co-crystal structure of WDR91 with XS381295 showed it binding to the same side pocket also in an enantioselective manner (Supplementary Fig. 11c and 12), even though this was not obviously apparent by E-ASMS. In the case of XS381295, its chlorophenyl ring fits into a hydrophobic pocket formed by the aliphatic side chains of L477, L465, L467, A459, as well as T547 and M550 (Supplementary Fig. 12). We suspect that the inability to detect enantioselective binding for XS381295 by chiral chromatography may be due to its rapid racemization.

*WD40 repeat containing protein 55 (WDR55)* is a 383-residue WDR protein that adopts a seven-bladed β-propeller fold and functions as a nucleolar protein involved in ribosomal biogenesis^28^. Although WDR55 is predicted to be a challenging target according to its DLID score, racemic compound XS381774 was enriched (Fig. 3g and 3h), with a subsequently confirmed apparent *K*_D_ of 11 μM, as measured by SPR, and displayed enantioselectivity (Supplementary Fig. 7). The enantioselective binding of XS381774 was of interest, because the enantiomers differed by the orientation of a single methyl group. The mechanism of enantioselective binding was revealed by co-crystallization of racemic XS381774 with WDR55 (Supplementary Fig. 11d and Table 4). The (*S*)-enantiomer of XS381774 preferentially co-crystallized in a hydrophobic side pocket of WDR55, and binding was mediated through hydrophobic interactions with the aliphatic side chains of residues I100, I108, V110, V122, L141, L135, and W153 (Fig. 3i). A methyl group which forms the chiral center points toward W153 and I108, enhancing hydrophobic interactions and that explained the enantioselective enrichment. To quantify the degree of selectivity, we separated and purified the individual enantiomers and evaluated their binding affinities to WDR55 using SPR. The (*S*)-enantiomer of XS381774 exhibited at least 5-fold higher binding affinity compared to the (*R*)-enantiomer, with *K*_D_ values of 5 μM and > 25 μM, respectively (Supplementary Fig. 7).

*Histone Acetyltransferase 1 (HAT1)* is an enzyme that acetylates histone lysine residues, a modification associated with a transcriptionally active chromatin state^30^. Despite two decades of efforts in both academia and industry, there are few, if any, small molecule ligands for HAT1. In our E-ASMS screen, XS380871 was selectively enriched by HAT1 (Fig. 3j). Interestingly, XS380871 contains two chiral centers, and two of its four stereoisomers, were selectively enriched by HAT1 (Fig. 3k). The binding of XS380871 to HAT1, was confirmed by SPR (*K*_D_ = 12 μM) as well as ^19^F NMR (Supplementary Fig. 8). To determine the molecular basis of binding, we incubated and co-crystallized the racemic mixture of XS380871 with HAT1 (Supplementary Fig. 11e and Supplementary Table 4). The co-structure revealed that the compound binds at the acetyl-Coenzyme A (acetyl-CoA) binding site (Fig. 3l). The chiral carbon (*S*) linked to the nitrophenyl moiety exhibited an enantioselective binding mode, with the nitrophenol substituent forming hydrogen bonds with the backbone nitrogen of G253 (via its nitro group) and the backbone nitrogen of G249 (via its hydroxyl group). There would however be enough space elsewhere in the structure to accommodate the (*R*)-enantiomer, albeit with potentially reduced affinity. In contrast, the fluorophenyl moiety at the second chiral carbon, bound to a hydrophobic subpocket lined by residues A275, V238, M241, and Y282; here the other enantiomer would be incompatible with binding due to severe steric clashes with the region around M241. This observation is consistent with the enantioselectivity data obtained from the E-ASMS analysis.

## Discussion

Despite the extensive efforts over the past decades^6^, approximately 80% of all human proteins still lack chemical ligands^5,7,8^. The slow pace of chemical ligand discovery is largely due to the intrinsic limitations of the current hit identification strategies. Here, we introduce the E-ASMS approach for high-throughput chemical ligand discovery: 1) The high sensitivity of the E-ASMS method in capturing weak-affinity hits ensures a high success rate, even with a relatively small chemical library (~8,000 compounds). This will reduce the time and financial cost for scalable screening. 2) Hit compounds may be able to be rapidly confirmed using orthogonal enantioselectivity information, bypassing time-consuming biophysical validations required for scalable and early-stage screening. The concept is that E-ASMS provides increased evidence of the specificity of compound binding not through use of an orthogonal assay with an orthogonal compound. Arguably, if a compound binds in an enantioselective manner in the screen at ~100 nM concentration, it may provide sufficient evidence of binding to obviate the need to test the compound at higher concentrations in an orthogonal biophysical assay. We envision that the E-ASMS approach will provide a convenient solution for scalable and proteome-wide chemical ligand discovery.

E-ASMS method may also be able to provide semi-quantitative information for identified hits. As shown in Supplementary Fig. 13, strong correlations were observed between the recovered MS signals of confirmed hits and their *K*_D_ values. This raises the possibility that the MS signals of E-ASMS hits can be used to generate large high-quality datasets that support artificial intelligence (AI)-based drug discovery.

Not all E-ASMS candidate binders could be confirmed using biophysical methods, such as SPR. Our limited data indicates this may be due to differences in their underlying biophysical principles: the SPR method can only detect hit compounds at concentrations above their *K*_D_, which requires compounds to be soluble at high concentrations whereas E-ASMS is insensitive to chemical concentrations. This discrepancy most apparent for weak E-ASMS hits, which may fail validation by SPR due to their low water solubility. Moving forward, we propose a general strategy of obtaining analogues of E-ASMS hits with greater predicted solubility to facilitate biophysical validation.

## Methods

### Reagents

Methanol (HPLC grade), ultrapure water (HPLC grade), acetonitrile (HPLC grade), DMSO, glycerol, and formic acid were purchased from Fisher Scientific (Ottawa, ON, CA). Ni-NTA magnetic agarose beads and Triton X-100 were obtained from Sigma-Aldrich (St. Louis, MO, USA). Tris base, sodium chloride (NaCl), and Tris(2-Carboxyethyl)phosphine (TCEP) were sourced from BioShop Canada Inc. (Burlington, ON, CA). Imidazole was provided by Bio Basic Canada Inc. (Markham, ON, CA). The chemical library was purchased from Enamine US Inc. (Monmouth Jct., NJ, USA) and ChemDiv (San Diego, CA, USA). Stock solutions of the chemicals was prepared in DMSO and stored at −20 °C in the dark until use.

### E-ASMS protein affinity selection experiments

The E-ASMS library of 8,210 chemicals was divided into 14 chemical pools by using a robotic system, with an average of ~600 chemicals in each pool. Each of the 14 chemical pools was incubated with purified recombinant His-tagged human proteins on 96-well plates, in the binding buffer (150 mM NaCl, 50 mM Tris-HCl, 0.1 mM TCEP, 0.5% (v/v) glycerol, 0.01% (v/v) Triton X-100, and pH 7.5). 5 µL of Ni-NTA magnetic beads was added to the binding buffer, adjusted to a final volume of 250 µL per sample. The final concentrations were set to 1 µM for the proteins and 100 nM for each compound. All E-ASMS screenings were performed in triplicate. Incubation was carried out on a rotary mixer at 4 °C for 30 min. After incubation, the 96-well plate was placed on a magnetic plate to separate the beads, and the incubation buffer was removed. The beads were then washed twice with 100 µL of washing buffer 1 (150 mM NaCl, 50 mM Tris-HCl, 0.1 mM TCEP, 0.5% glycerol, 0.01% Triton X-100, 5 mM imidazole, and pH 7.5), followed by one wash with 100 µL of washing buffer 2 (150 mM NaCl, 50 mM Tris-HCl, 5 mM imidazole, and pH 7.5). The beads and washing buffer 2 were transferred to a new plate. After removing the washing buffer 2, 120 µL of methanol was added to each well to denature proteins and release chemical ligands. The methanol extracts separated from magnetic beads were directly subjected to LC-MS analysis. In parallel, the proteins were also subjected to SDS-PAGE to confirm the presence of target proteins on the beads.

### Identification of putative hits using LC-MS

The E-ASMS extracts were analyzed using an Orbitrap Exploris 240 mass spectrometer equipped with a Vanquish UHPLC system (Thermo Fisher Scientific, CA, USA). Chromatographic separation was conducted on an Accucore Vanquish C_18_ reverse-phase column (50 mm × 2.1 mm × 3 µm) at a flow rate of 0.3 mL/min. The injection volume was 1 µL. Ultrapure water with 0.1% formic acid (A) and methanol with 0.1% formic acid (B) were used as the two mobile phases. Gradient elution started with 5% B, increased to 80% at 1.5 min, then reached to 100% at 3 min, held for 1.1 min, and finally returned to 5% B over 1.5 min. The column temperature was maintained at 40 °C, and the sample compartment at 7 °C. Data acquisition was performed in full MS^1^ scan mode (150-520 *m/z*) with a resolution of R = 60,000, in both positive and negative ionization modes. The detailed instrumental parameters of LC-MS system are provided in Table S5.

### Chiral analysis of putative hits

*1) Building the chiral separation database:* the accurate prediction of suitable chiral separation conditions for resolving the enantiomers of a given compound remains a challenge^31,32^. To address this, we decided to establish the chiral separation database for the E-ASMS chemical library, under eight conditions (4 columns × 2 LC gradient methods, see the details below). Each of the 14 chemical pools were individually injected under eight conditions to build the database.

*2) Eight chiral separation conditions:* four columns from DAICEL Chemical Industries, LTD. were employed: CHIRALPAK IA (250 × 4.6 mm), CHIRALPAK IBN (250 × 2.1 mm), CHIRALPAK IC (250 × 2.1 mm), and CHIRALPAK IG (250 × 2.1 mm). These columns were selected because they have been demonstrated to achieve sufficient separations for the biggest number of chiral compounds^31^.

Two LC gradient methods were employed for each column. In **method 1**, acetonitrile with 0.1% formic acid (A) and methanol with 0.1% formic acid (B) were used as mobile phases. Gradient elution started at 40% B, held for 1 min, then increased to 80% at 3 min, reached 100% at 8 min, held for 1 min, followed by a return to 40% over 2 min, and held for another 1 min. The flow rate was set at 0.3 mL/min for columns IBN, IC, IG, and at 1.0 mL/min for column IA.

In **method 2**, H_2_O with 0.1% formic acid (A) and methanol with 0.1% formic acid (B) were used as mobile phases. Gradient elution began with 40% B, increased to 80% at 6 min, reached 100% at 18 min, held for 6 min, and then return to 40% over 6 min. The flow rate was 0.2 mL/min for columns IBN, IC, IG, and 1.0 mL/min for column IA. The column temperature was maintained at 40 °C, and the sample compartment was kept at 7 °C. The injection volume was 1 µL.

*3) Chiral detection of putative hits under optimal conditions:* Once putative hits were detected from initial E-ASMS screening, the optimal separation condition for the hit compounds was selected by searching against the chiral separation database established in step 2) above. The same E-ASMS extracts were then analyzed under the optimal conditions. To exclude potential interferences, enantiomers were further confirmed by matching MS^2^ spectra.

### Automatic data processing

A total of 1,302 E-ASMS screening samples (31 proteins × 3 replicates × 14 pools = 1,302) were completed in the current study, leading to 145 GB of raw mass spectrometry data. We have established an automatic data processing workflow to detect E-ASMS hits.

*1) Building the MS database for the E-ASMS library*: To build the database, each of the 14 chemical pools was subjected to LC-MS analysis. Raw mass spectrometry files were converted to the mzXML format. Peak features from each chemical pool were identified using the ‘XCMS’ R package with a mass tolerance of 2.5 ppm^29^. Only the peak features with peak an intensity > 10^5^ and abundances at least 10 times higher than methanol were considered as library compounds and retained for subsequent analysis. Isotopic peaks and adducts were excluded by matching chromatographic peaks and theoretical mass difference. Detected peak features were then matched to the E-ASMS library with a mass tolerance of 3 ppm. Potential inferences and misassigned compounds were further excluded via manual inspections. Then, a MS database of 8,210 E-ASMS library compounds were established, with *m/z*, retention time and SMILES information recorded.
*2) Detecting compounds from E-ASMS features by matching to the MS database*: To detect library compounds from the E-ASMS features, we matched each of the 8,210 E-ASMS compounds against each E-ASMS samples. A mass tolerance of 3 ppm, and a retention time matching window of 0.5 min were used. A final data matrix with 8,210 rows (corresponding to library compounds) and 24 columns (corresponding to E-ASMS samples from each batch) was created.
*3) Detecting putative E-ASMS hits*: To detect putative E-ASMS hits, we calculated the enrichment fold and *p* values of each library compound against each protein by using seven other proteins from the same batch as the control (equation 1).

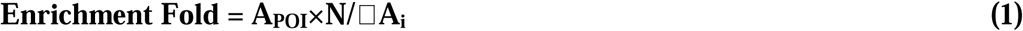

A_POI_ represents the peak intensity of the putative hit compound enriched by the protein of interest; N represents the number of background proteins from the same batch (N = 7); A_i_ represents the peak intensity of the same hit compound enriched by the i^th^ protein from the same batch.

Eventually, only the compounds with enrichment fold > 5, and *p* value < 0.05 were considered as putative hit compounds. Each putative hit was further manually inspected by matching to the chemical library, to exclude potential interferences misassigned by the algorithms.

### Quality Assurance and Quality Control (QA/QC) of E-ASMS

To assure the high-quality data for scalable E-ASMS screening, we conducted stringent QA/QC from both protein and chemical perspectives.

### Protein QA/QC

To assure the selected proteins were properly folded and compatible with our E-ASMS workflow, we used SDS-PAGE to semi-quantitatively assess the recovery of each protein through our E-ASMS procedure. In brief, SDS-PAGE analysis was conducted for each protein before and after E-ASMS. The band intensity of each protein was compared, and those proteins with low recoveries were excluded for E-ASMS analysis. The SDS-PAGE results for all proteins are provided in Supplementary Fig. 14.

### Chemical QA/QC

During LC-MS analysis, the E-ASMS chiral library compounds were injected every batch to ensure the retention time, *m/z*, and intensity are not shifting for database matching. All the chiral compounds were detected by the LC-MS system. A QC standard of small subset (~600 chemicals) was injected after every 12 samples to monitor instrument stability and LC (chiral) separations. The instrument response deviation for analytes in the QC standard remained below 10% throughout the analysis. A solvent blank was injected after every 12 samples to monitor and prevent carryover.

### Proteins used for E-ASMS and SPR

Protein expression and purification summaries are detailed in Supplementary Table 6.

### Crystallization and structure determination

Crystallization of E-ASMS hits with their respective targets are summarized in Supplementary Table 7.

### Surface plasmon resonance

SPR hit confirmation and validations were performed using a Biacore 8K instrument at 20 °C. The biotinylated respective protein constructs were immobilized on the active flow cells of a streptavidin-coated SA sensor chip after initial conditioning of both reference and active flow cells with 50 mM NaOH for 3 × 60 s with a flow rate of 10 μL/min. Each protein solution (30-150 μg/mL) was injected through the respective active flow cells for 60-660 s with a flow rate of 5 μL/min to obtain a protein immobilization level estimated to yield ~30 RU of signal at 100% compound binding. Equilibration was performed after protein immobilization by flowing running buffer (50 mM HEPES, pH 7.5, 150 mM NaCl, 0.001% Tween 20, 0.2% PEG3350, 0.5 mM TCEP, and 3% DMSO) over the flow cells with a flow rate of 50 μL/min until a stable baseline was observed. Initial start-up cycles, blank cycles and wash (50% DMSO wash to flush the needles) steps were included in the SPR compound analysis methods. The compound injections were performed over the reference and active flow cells using multi-cycle kinetics at a flow rate of 40 μL/min with a 55 s association time and a 120 s dissociation time for dose-response titration. The stock compound concentration series was performed in 100% DMSO with 2-3-fold serial dilution and prepared the samples using the running buffer by maintaining final DMSO concentration of 3% (v/v) across the tested concentration range. Initially, the compounds were titrated between 0-45 µM compound concentrations and later followed up between 0-15 µM, 0-30 µM, 0-45 µM, and 0-90 µM compound concentrations based on the affinity and solubility of the compounds. Solvent correction cycles were also included across each run to adjust high bulk responses from the solvent. Double referencing of the data was introduced by subtraction of the reference flow cell and the respective zero compound concentration cycles. Affinity fitting was performed by applying a 1:1 equilibrium binding model to the data using Biacore Intelligent Analysis tool provided with the Biacore Insight Evaluation software.

### 19F NMR studies

The binding of XS380871 to HAT1 was confirmed using ^19^F NMR by looking for the broadening and/or shifting of its ^19^F resonance upon the addition of HAT1. Spectra of the compound were acquired on a Bruker Avance III spectrometer operating at 600 MHz, equipped with a QCI probe at 293K, and collected at 20 µM, with and without the presence of HAT1 (~34 µM, buffered in 50 mM Tris 7.0, 200 mM NaCl, 4 mM DTT). TFA (100 µM) was added as an internal reference. 1k transients were acquired over a sweep width of 150 ppm; an exponential window function (LB = 5 Hz) was applied prior to Fourier transformation.

### Statistics

All statistical analyses were conducted using GraphPad PRISM (GraphPad Software, Inc., La Jolla, CA). Data are presented as mean ± standard deviation. Statistical significance was determined using Student’s *t*-test, with a significance threshold of *p* < 0.05.

## Data availability

Atomic coordinates and structure factors for all crystal structures have been deposited in the Protein Data bank under the accession codes: 9EJO, 9EJP, 9EJQ, 9EKP, 9MJG.

## Code availability

The source code and demo datasets can be found via GitHub at https://github.com/huiUofT/E-ASMS.

## Acknowledgements

We would like to thank the advice and discussions on method development from Dafydd Owen (Pfizer), Judith Guenther (Bayer), Scott Johnson (Bristol Myers Squib), Oliver Kraemer (Boehringer Ingelheim), Julian Schmid (Merck), Juncai Meng (Janssen), Lawrence Szewczuk (Janssen), Xidong Feng (Pfizer), Wenyi Hua (Pfizer), Anja Giese (Bayer), Matteo Aldeghi (Bayer), Nidhi Arora (Takeda), James Kiefer (Genentech), Ingo Hartung (Merck), and Matthew Troutman (Pfizer). The Structural Genomics Consortium is a registered charity (no. 1097737) that receives funds from Bayer AG, Boehringer Ingelheim, Bristol Myers Squibb, Genentech, EU/EFPIA/OICR/McGill/KTH/Diamond Innovative Medicines Initiative 2 Joint Undertaking [EUbOPEN grant 875510], Janssen, Pfizer, and Takeda. Research was supported by a Natural Science and Engineering Research Council of Canada (NSERC) Discovery Grant, Ontario Early Researcher Award, and the National Institute of Health (NIH) (U54AG065187). The authors acknowledge the support of instrumentation grants from the Canada Foundation for Innovation, the Ontario Research Fund, and an NSERC Research Tools and Instrument Grant. A.C.J, S.M., and S.K. are also grateful for support by the German Cancer Aid grant TACTIC. TSB acknowledges startup funding provided by the Emory University School of Medicine and the Winship Cancer Institute. AME holds the Temerty Nexus Chair of Health Technology and Innovation at the University of Toronto.

## Author contributions

XW, JS, DY, AB, SA, PJB, VS, LH, HP, AME: E-ASMS method development

JS, XW: E-ASMS screening

HZ, PG, EG, MK, SRG, LKR, DM, MS, CL: Protein production and QC

ShA, MK, FL, SRG, MS: SPR, DSF confirmation

SH: F-NMR characterization

HZ, UHC, AD: X-ray crystallography

MS, CVS: Druggability assessment

SA, VS, PJB: Project management

WWY, RH, TSB, SM, ACJ, SK, LH, HP, AME, CHA: Supervision, review and editing

XW, JS, LH, HP: Writing, reviewing and editing with input from all

## Competing interests

The authors declare no competing interests.

## Notes

### Competing Interest Statement

The authors have declared no competing interest.

### Summary of Updates

The name of the approach was changed from EAS-MS to E-ASMS.

