## Supplementary Materials for "Enantioselective Protein Affinity Selection Mass Spectrometry (E-ASMS)"

P. 8 – Supplementary Fig. 4 | Distribution of E-ASMS hits across the chemical space.

P. 9 – Supplementary Fig. 5 | Representative SPR sensorgrams and the response vs concentration plots for the DDB1 specific hit.

P. 10 – Supplementary Fig. 6 | Representative SPR sensorgrams and the response vs concentration plots for the four WDR91 specific hits.

P. 11 – Supplementary Fig. 7 | Representative SPR sensorgrams and the response vs concentration plots for the WDR55 specific hit.

P. 12 – Supplementary Fig. 8 | Representative SPR sensorgram and the steady-state response vs concentration plot for the HAT1 specific hit XS380871.

P. 13 – Supplementary Fig. 9 | Representative SPR sensorgrams and the response vs concentration plots for the SKP1, DCAF1, DCAF7, FBXW7, SKP1-FBXO22, AASS(LKR), and AASS(SDH) hits.

P. 14 – Supplementary Fig. 10 | ECD analysis of two enantiomers of XS381952 binding to DDB1.

P. 15 – Supplementary Fig. 11 | The mFo-dFc electron density omit-maps for the E-ASMS hits in the crystal structures.

P. 16 – Supplementary Fig. 12 | Co-crystal structure of WDR91 in complex with XS381295.

P. 17 – Supplementary Fig. 13 | Strong correlations between hit MS signals and *K*_D_ values.

P. 18 – Supplementary Fig. 14 | Uncropped SDS-PAGE images.

P. 19-20 – Supplementary Table 4. Data collection and refinement statistics.

P. 21 – Supplementary Table 5. Detailed instrumental parameters of LC-MS.

Supplementary Tables 1-3, 6, and 7 are provided as separate Excel sheets.

Supplementary Table 1. Chiral compound library.

Supplementary Table 2. Positive ligands.

Supplementary Table 3. Candidate binders.

Supplementary Table 6. Purification methods.

Supplementary Table 7. Crystallization information.

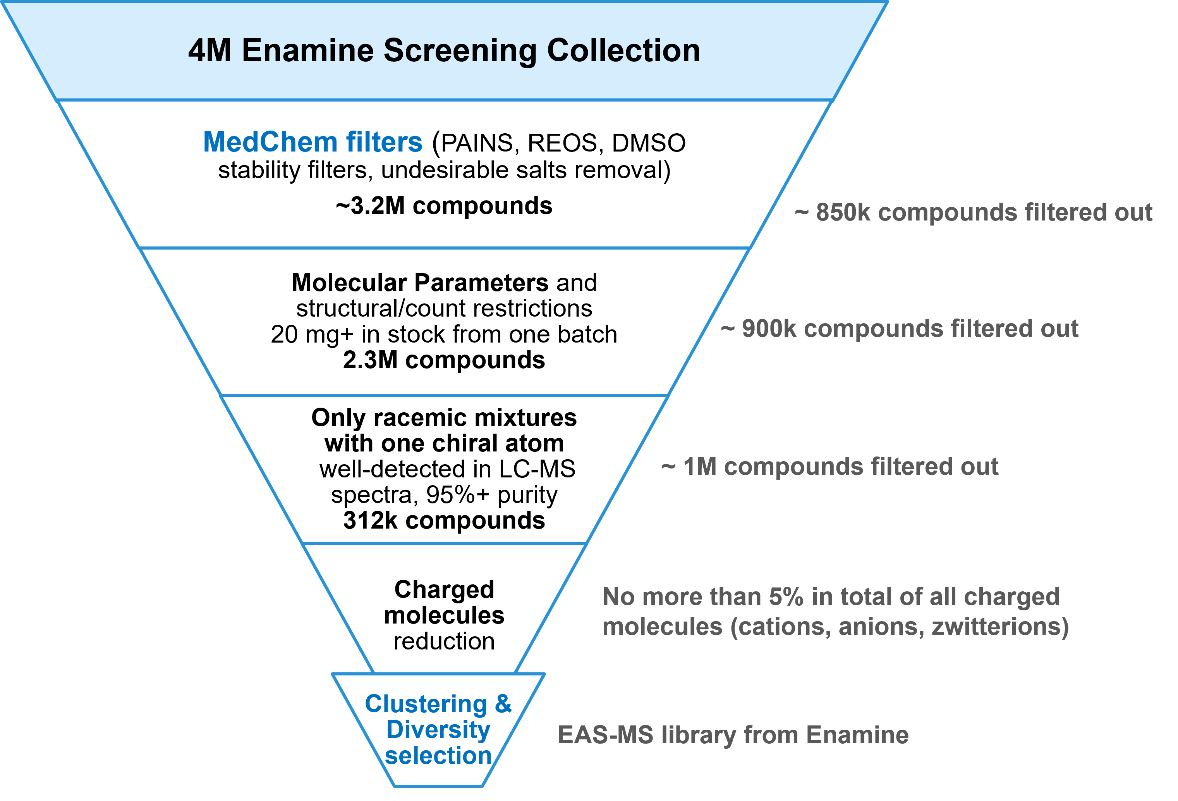

Supplementary Fig. 1 | Strategy for E-ASMS chemical library selection.

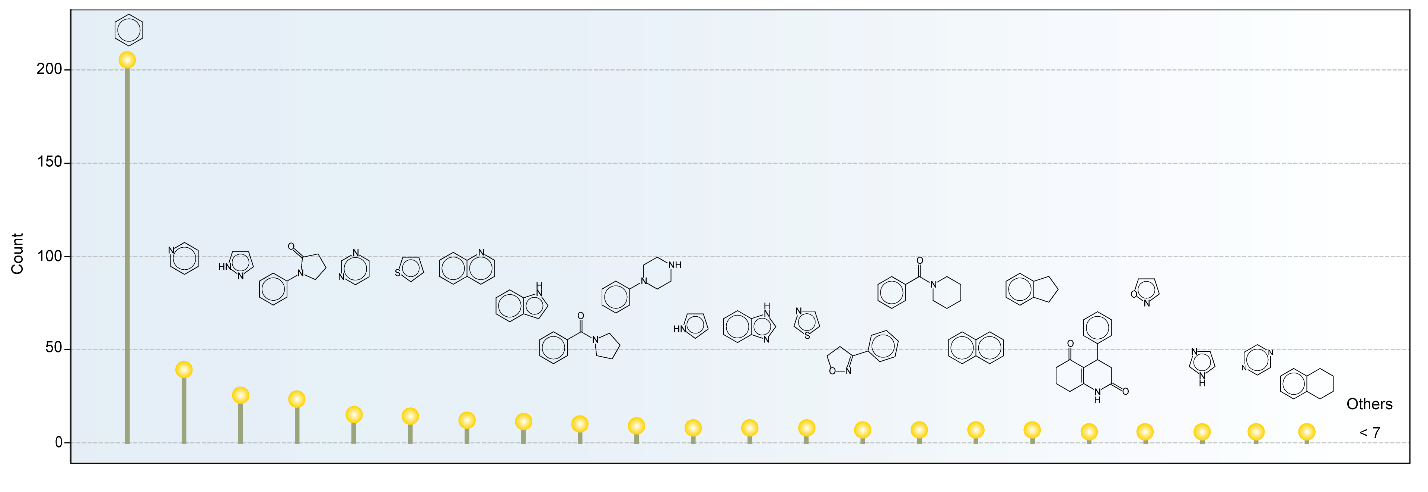

Supplementary Fig. 2 | Top 20 scaffolds in the E-ASMS compound library. The E-ASMS library comprises a total of 7,307 distinct scaffolds.

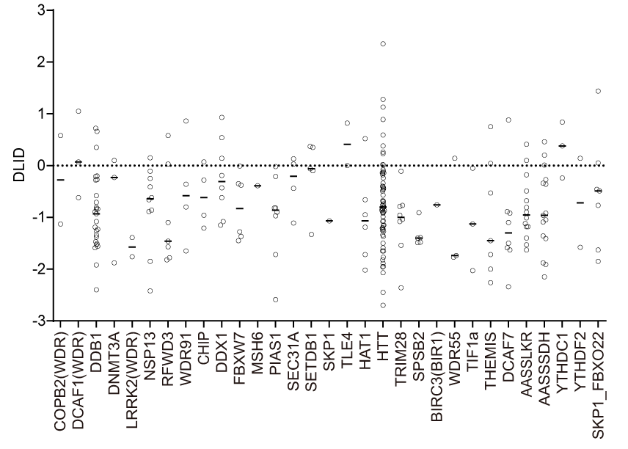

Supplementary Fig. 3 | Predicted ligandability of 31 selected proteins. Drug-like density (DLID) of each protein was predicted. The dots from each protein represent the predicted DLID score for each pocket in that protein. To define the ligandability of proteins, we used the maximal DLID score for the pocket from each protein. For instance, the maximal DLID score for HTT is 2.35 for the top drug-like pockets compared to other pockets, due to its large molecular size and surface. Proteins with low, medium and high ligandability were defined as having maximal DLID scores < 0.5, 0.5-1, and > 1, respectively.

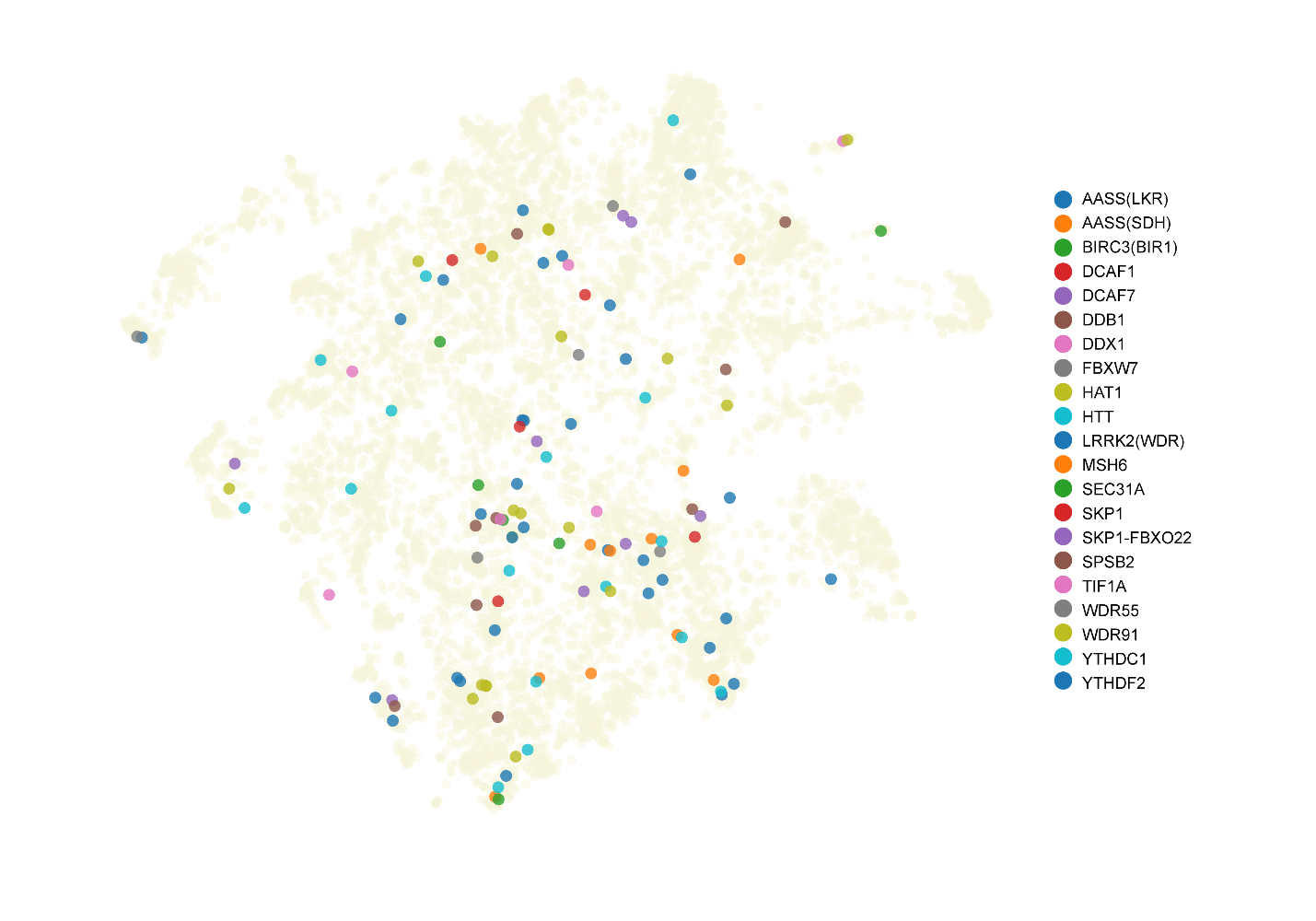

Supplementary Fig. 4 | Distribution of E-ASMS hits across the chemical space. UMAP visualization based on SMILES descriptor highlights the chemical distribution of identified hits.

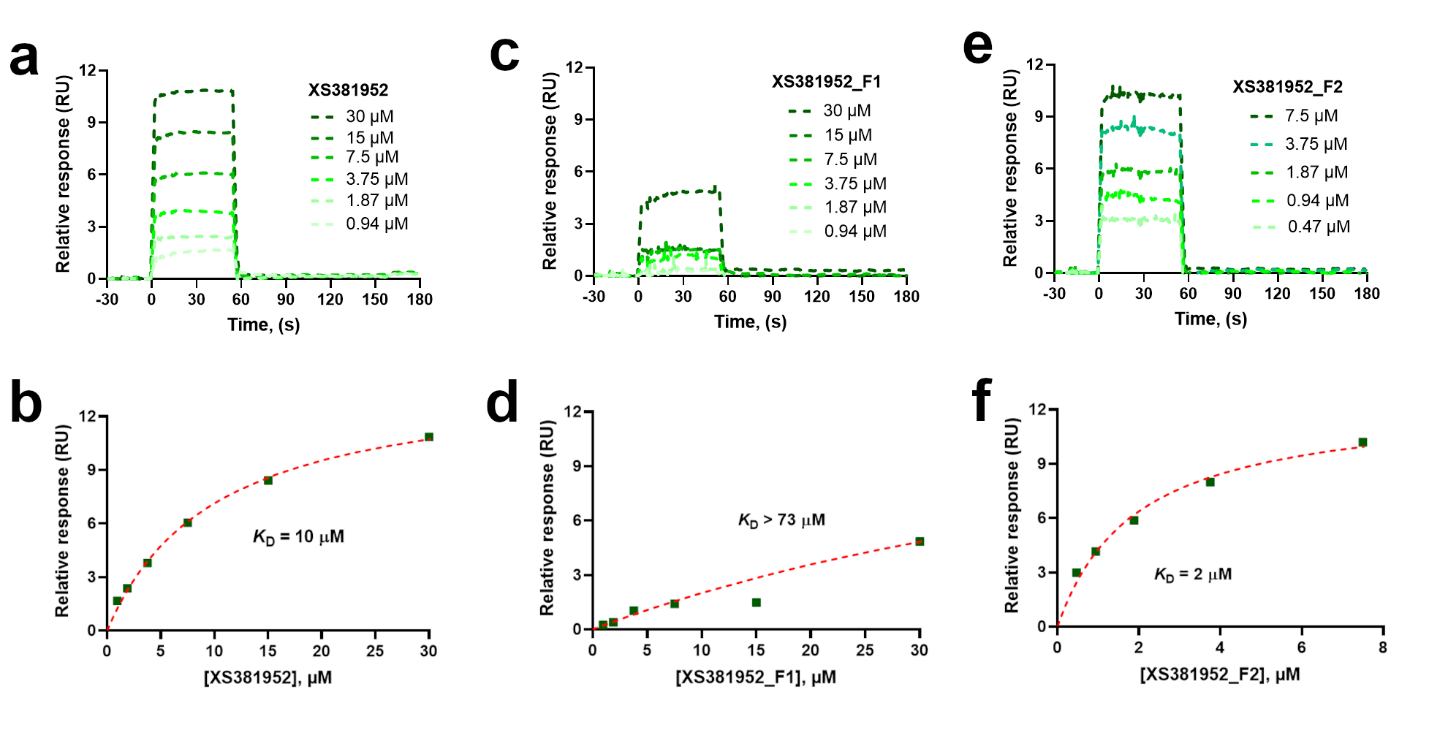

Supplementary Fig. 5 | Representative SPR sensorgrams (a, c, e) and the response vs concentration plots (b, d, f) for the DDB1 specific hit XS381952 (a, b) and its enantiomers XS381952_F1 (c, d) and XS381952_F2 (e, f) after separation, fraction 1 (100% pure) and fraction 2 (85% pure), respectively. The response at equilibrium for each concentration was plotted against the compound concentration and affinity fitted by applying 1:1 equilibrium binding model to extract the *K*_D_ of the interaction. The compounds were tested in duplicates. Data from one replicate is shown here for clarity.

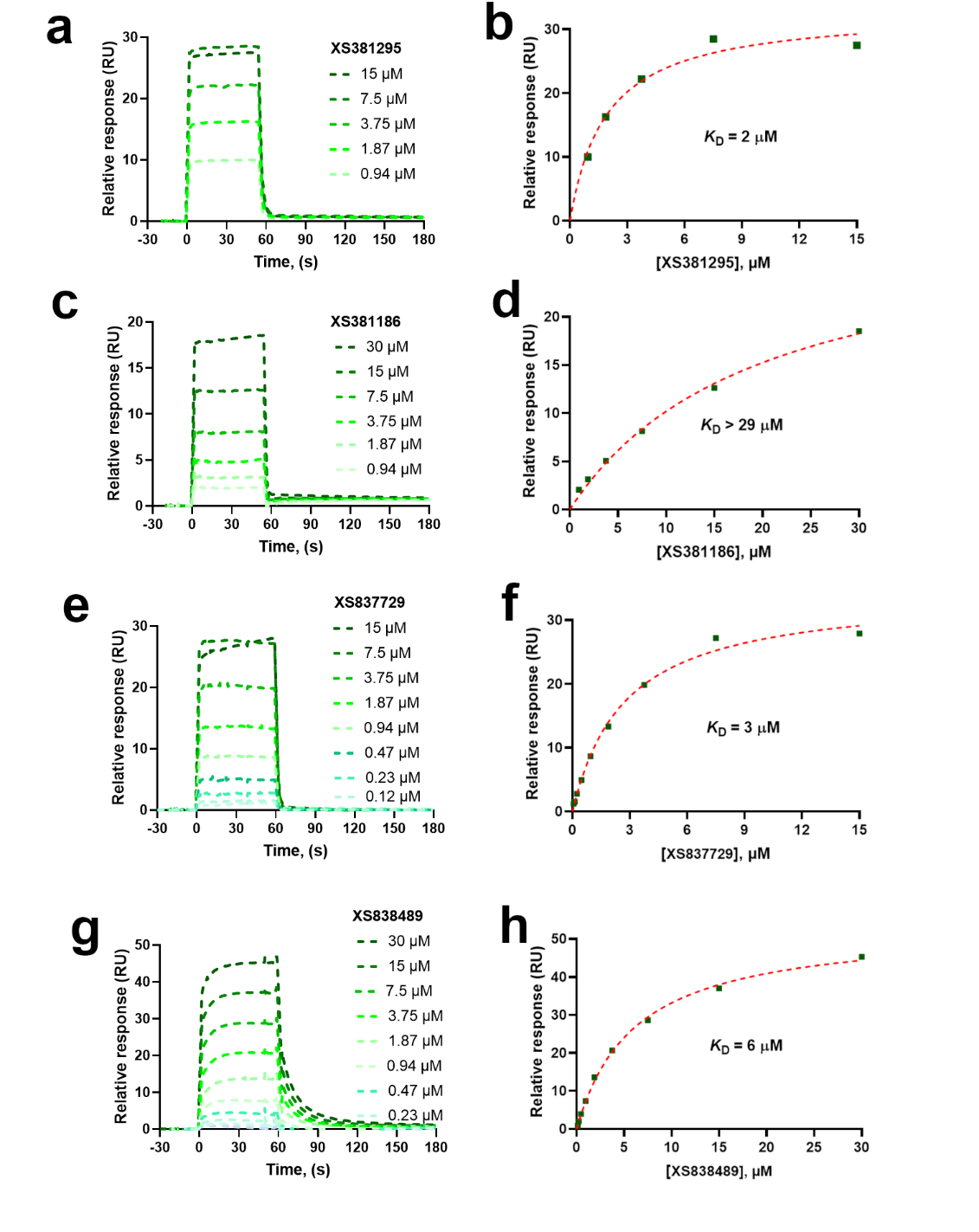

Supplementary Fig. 6 | Representative SPR sensorgrams (a, c, e, and g) and the response vs concentration plots (b, d, f, and h) for the four WDR91 specific hits: XS381295a (a, b), XS381186a (c, d), XS3837729 (e, f), and XS838489 (g, h). The response at equilibrium for each concentration was plotted against the compound concentration and affinity fitted by applying 1:1 equilibrium binding model to extract the *K*_D_ of the interaction. The compounds were tested in duplicates. Data from one replicate is shown here for clarity.

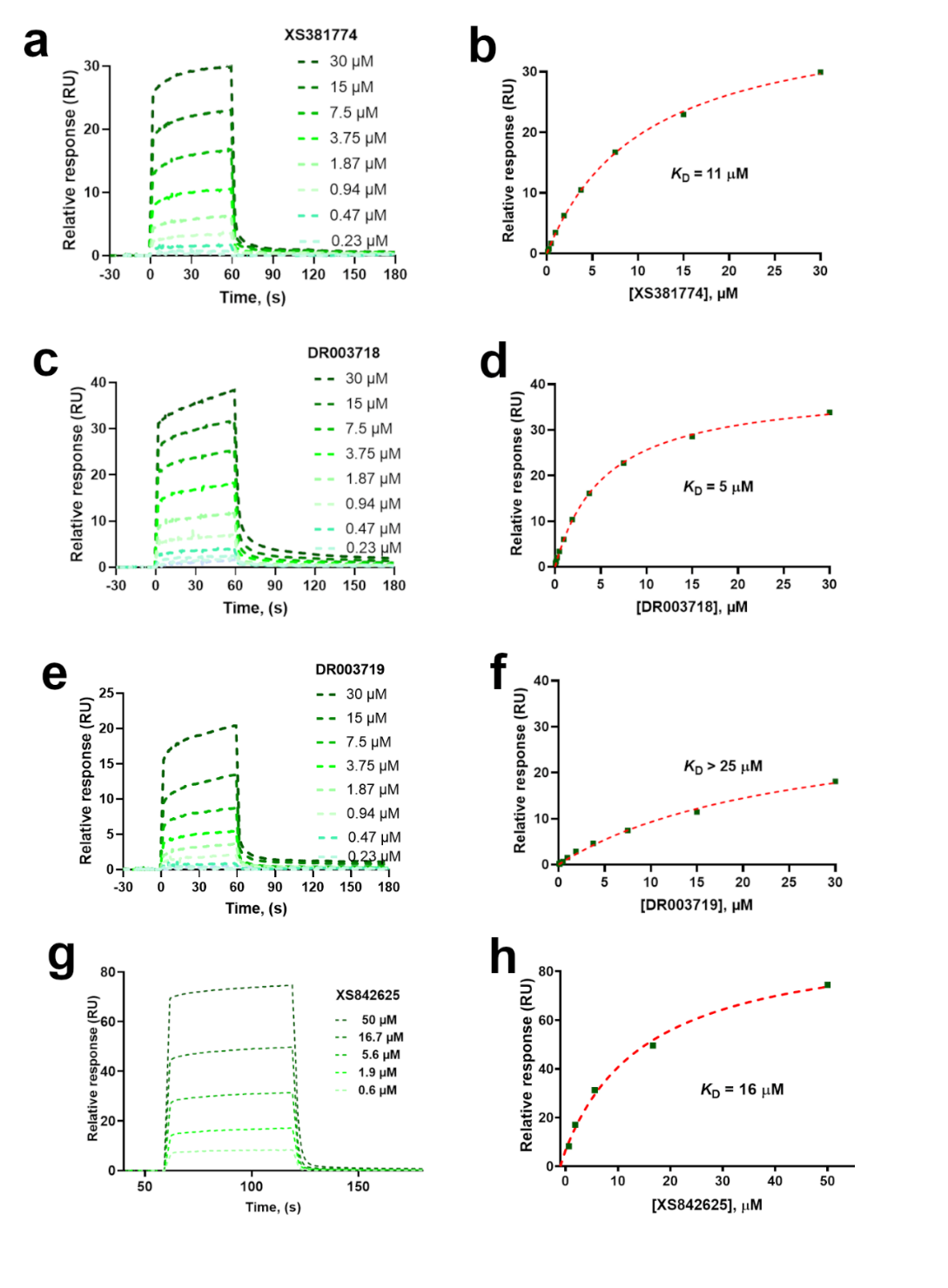

Supplementary Fig. 7 | Representative SPR sensorgrams (a, c, e, and e) and the response vs concentration plots (b, d, f, and h) for the WDR55 specific hit XS381774 (a, b) and its separated enantiomers DR003718 (c, d) and DR003719 (e, f), as well as XS842625 (g, h), respectively. The response at equilibrium for each concentration were plotted against the compound concentration and affinity fitted by applying 1:1 equilibrium binding model to calculate the *K*_D_ of the interaction. The compounds were tested in duplicates. Data from one replicate is shown here for clarity.

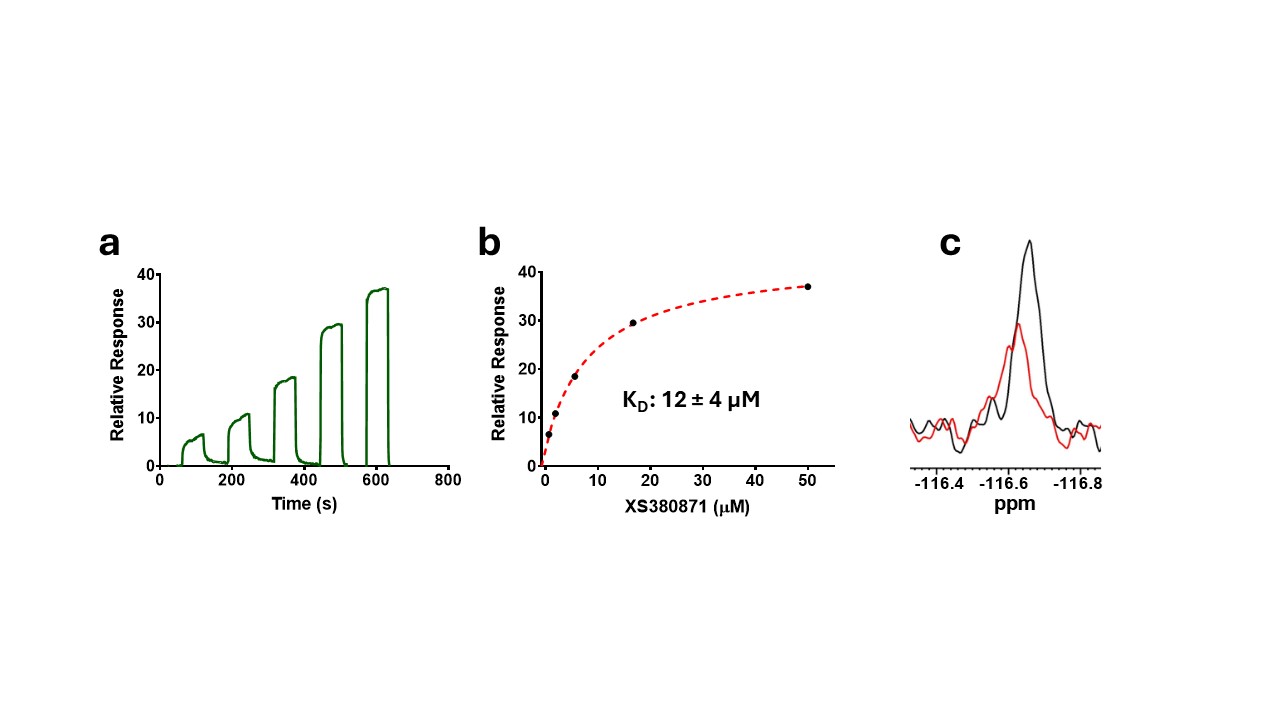

Supplementary Fig. 8 | Representative SPR sensorgram (a) and the steady-state response vs concentration plot (b) for the HAT1 specific hit XS380871. The response at equilibrium for each concentration were plotted against the compound concentration and affinity fitted by applying 1:1 equilibrium binding model (red dashed line), yielding a *K*_D_ value of 12 ± 4 µM. The experiments were performed in triplicate (n = 3). Data from one replicate is shown here for clarity. (c) Overlay of ^19^F spectra of 20 µM XS380871 alone (black), and in the presence of ~34 µM of HAT1 (red).

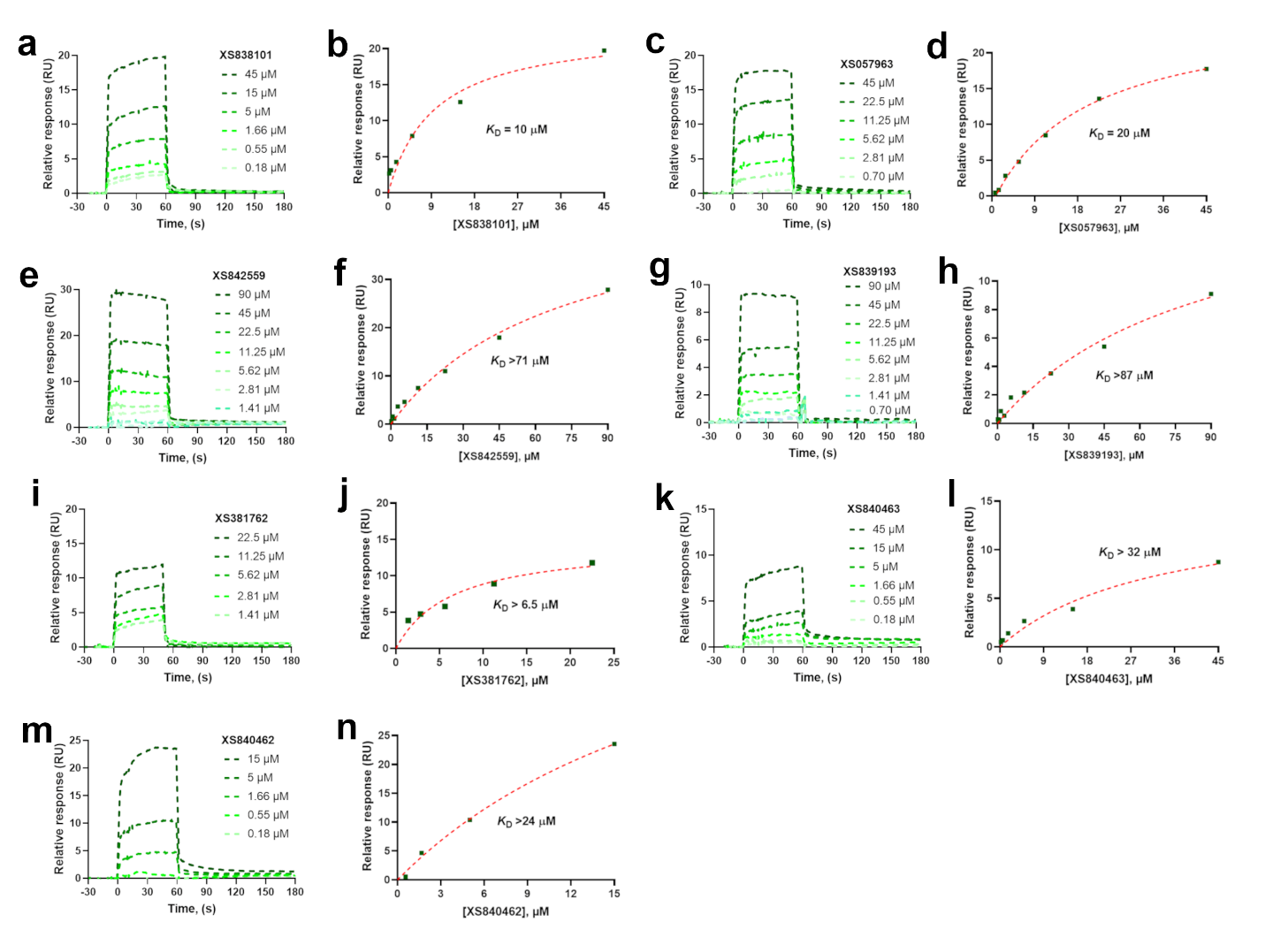

Supplementary Fig. 9 | Representative SPR sensorgrams (a, c, e, g, i, k, and m) and the response vs concentration plots (b, d, f, h, j, l, and n) for SKP1 (a, b), DCAF1 (c, d), DCAF7 (e, f), FBXW7 (g, h), SKP1-FBXO22 (i, j), AASS(LKR) (k, l), and AASS(SDH) (m, n) hits. The responses at equilibrium for each concentration were plotted against the compound concentration and affinity fitted by applying 1:1 equilibrium binding model to extract the *K*_D_ of the interaction. Where the *K*_D_ values are less/not reliable due to either sub stoichiometric binding or suboptimal concentration ranges for the dose-response titration were annotated by (>) sign. The compounds were tested in duplicates. Data from one replicate is shown here for clarity.

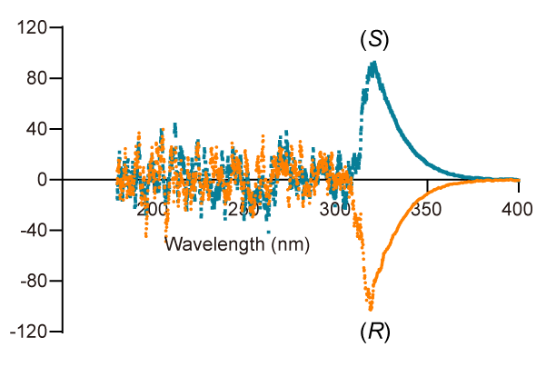

Supplementary Fig. 10 | ECD analysis of two enantiomers of XS381952 binding to DDB1. The two enantiomers were purified through the CHIRALPAK IA column.

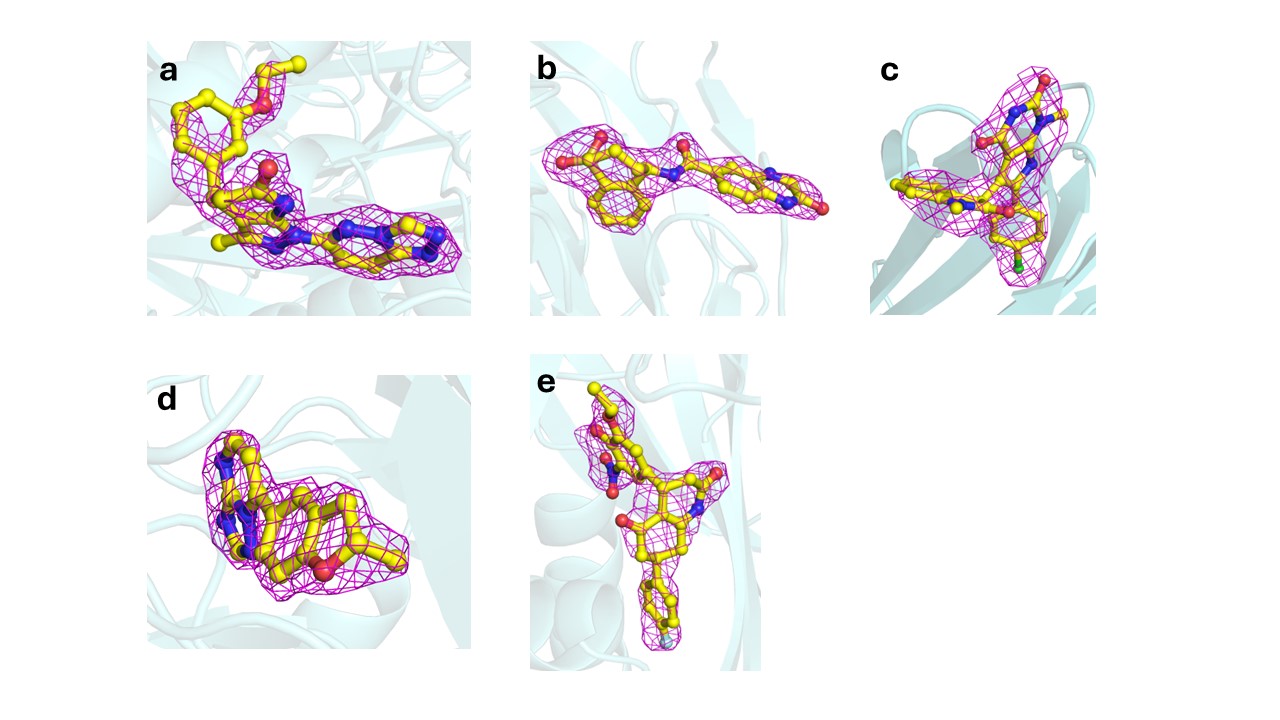

Supplementary Fig. 11 | The mFo-dFc electron density omit-maps for the E-ASMS hits (shown in yellow sticks) in the crystal structures of (a) DDB1_XS381952, (b) WDR91_XS838489, (c) WDR91_XS381295, (d) WDR55_XS381774, and (e) HAT1_XS380871 displayed as magenta meshes and contoured at 3σ.

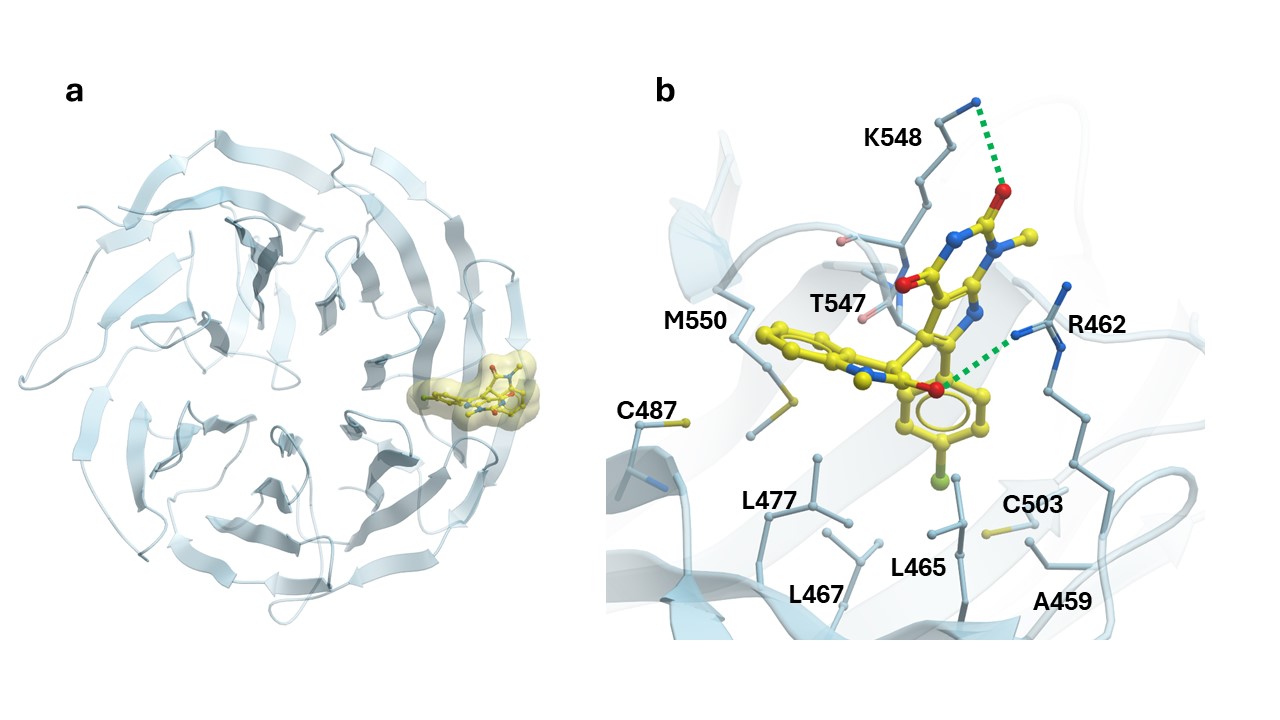

Supplementary Fig. 12 | Co-crystal structure of WDR91 in complex with XS381295. (a) Cartoon representation of WDR91 (cyan) bound to ligand (yellow). (b) Close-up view of the XS381295 binding site. Hydrogen bonds between the protein and the ligand are shown as green dashed lines.

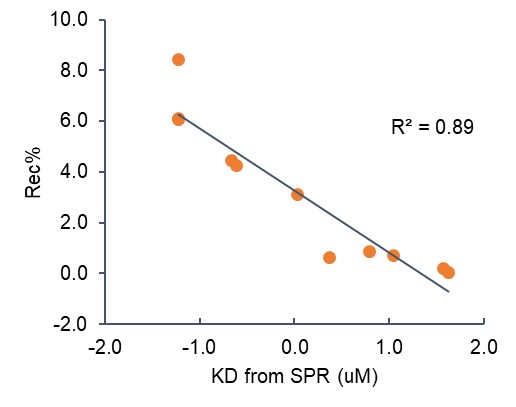

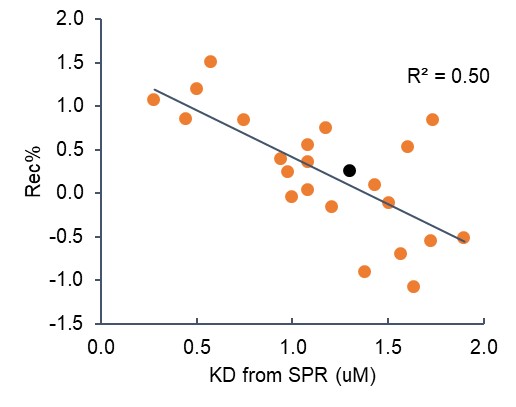

Supplementary Fig. 13 | Strong correlations between hit MS signals and *K*_D_ values of confirmed hits for single protein (WDR5, left) and all proteins (right).

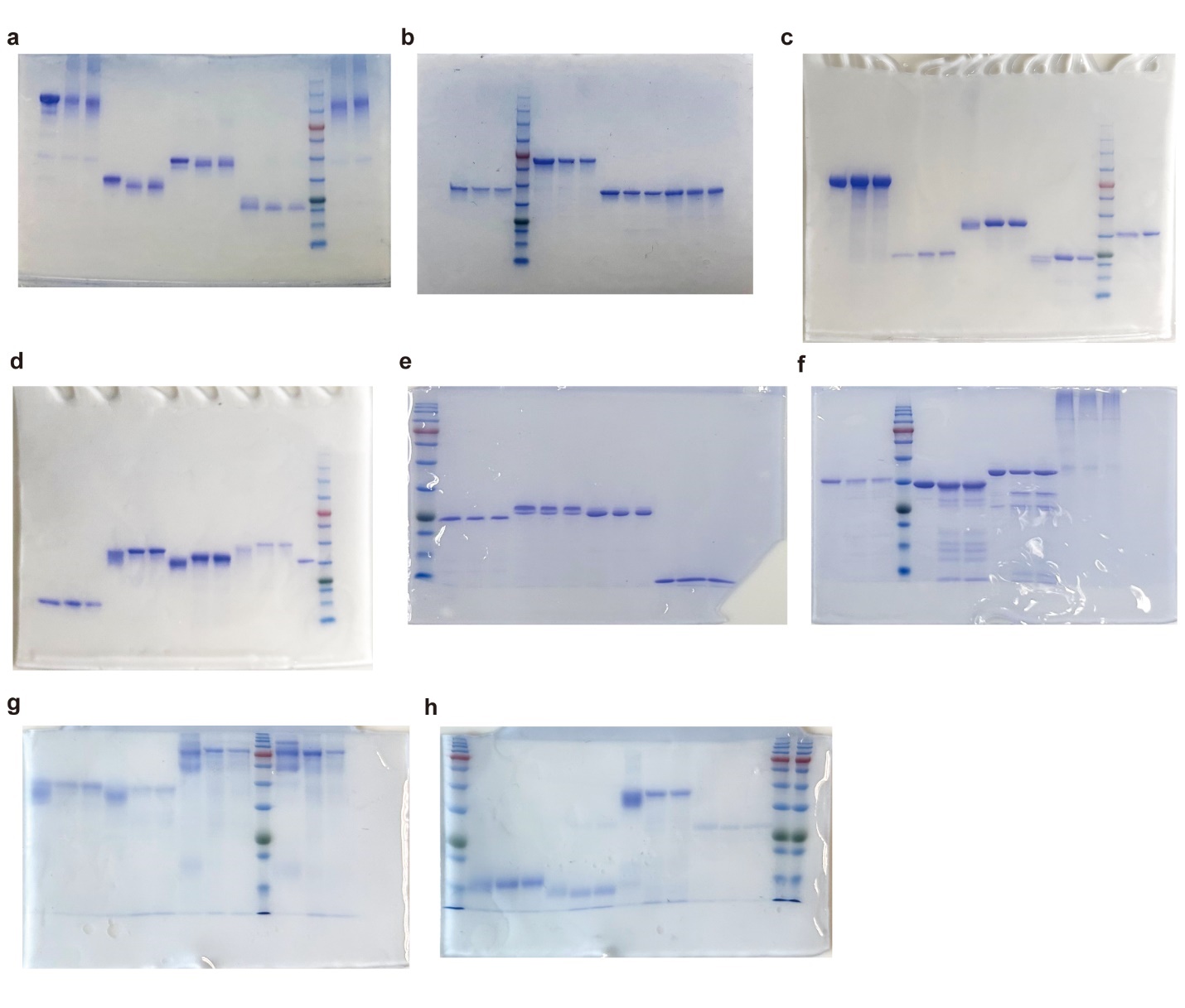

**Supplementary Fig. 14 |** **Uncropped SDS-PAGE images.** Each protein is shown in three lanes: total protein amount and two E-ASMS samples. (**a**) DDB1, COPB2(WDR), DCAF1(WDR), and DNMT3A. (**b**) LRRK2(WDR), NSP13, WDR91, and RFWD3. (**c**) DDX1, SETDB1, SEC31A, and SKP1. (**d**) MSH6, FBXW7, CHIP, and PIAS1. (**e**) TIF1A, TRIM28, SPSB2, and BIRC3(BIR1). (**f**) WDR55, TLE4, HAT1, and HTT. (**g**) THEMIS, DCAF7, AASS(LKR), and AASS(SDH). (**h**) YTHDC1, YTHDF2, and SKP1-FBXO22.

Supplementary Table 4. Data collection and refinement statistics.

|  | WDR91_XS381295 | WDR91_XS838489 | DDB1_XS381952 |
| --- | --- | --- | --- |
| PDB code | 9EJO | 9EJP | 9EJQ |
| **Data collection** |  |  |  |
| Space group | C222_1_ | C222_1_ | P2_1_2_1_2_1_ |
| Cell dimensions |  |  |  |
| *a*, *b*, *c* (Å) | 78.63, 132.38, 119.60 | 77.61, 132.55, 119.80 | 62.64, 124.70, 167.72 |
| α, β, γ (°) | 90.0, 90.0, 90.0 | 90.0, 90.0, 90.0 | 90.0, 90.0, 90.0 |
| Resolution (Å) | 50.0-2.40 (2.44-2.40) * | 50.0-2.22 (2.26-2.22) * | 50.0-1.87(1.90-1.87) * |
| R_sym_ or R_merge_ | 0.086 (0.966) | 0.062 (0.904) | 0.068 (0.940) |
| CC1/2 | 0.999 (0.716) | 1.001 (0.802) | 0.999 (0.666) |
| I / σI | 18.9 (1.9) | 31.0 (2.7) | 18.8 (1.9) |
| Completeness (%) | 99.9 (100.0) | 100.0 (100.0) | 91.2 (94.1) |
| Redundancy | 6.0 (6.0) | 10.8 (10.8) | 4.3 (4.7) |
| **Refinement** |  |  |  |
| Resolution (Å) | 27.70-2.40 (2.46-2.40) | 44.67-2.22 (2.28-2.22) | 25.10-1.87 (1.92-1.87) |
| No. reflections | 23892 | 29329 | 94637 |
| *R*_work_ / *R*_free_(%) | 19.9/23.8 | 20.3/24.5 | 16.4/22.0 |
| No. atoms | 2650 | 2682 | 9508 |
| Protein | 2535 | 2561 | 8739 |
| Ligand/ion | 30 | 27 | 29 |
| Water | 84 | 94 | 624 |
| *B*-factors | 48.0 | 48.5 | 31.5 |
| Protein | 48.1 | 48.5 | 30.8 |
| Ligand/ion | 55.5 | 57.6 | 45.2 |
| Water | 41.2 | 46.5 | 37.9 |
| R.m.s. deviations |  |  |  |
| Bond lengths (Å) | 0.006 | 0.005 | 0.007 |
| Bond angles (°) | 1.295 | 1.342 | 1.282 |

*Values in parentheses are for highest-resolution shell.

**Supplementary Table 4. Data collection and refinement statistics (continued).**

|  | WDR55_XS381774 | HAT1_XS380871 |
| --- | --- | --- |
| PDB code | 9EKP | 9MJG |
| **Data collection** |  |  |
| Space group | P2_1_ | P1 |
| Cell dimensions |  |  |
| *a*, *b*, *c* (Å) | 78.31, 58.41, 87.22 | 78.89, 87.84, 118.38 |
| α, β, γ (°) | 90.0, 96.3, 90.0 | 84.38, 80.07, 76.76 |
| Resolution (Å) | 34.81-1.95 (2.00-1.95) * | 50.0-2.58 (2.62-2.58) * |
| R_sym_ or R_merge_ | 0.082 (0.685) | 0.127 (0.653) |
| CC1/2 | 0.998 (0.774) | 0.984 (0.602) |
| I / σI | 14.1 (2.4) | 8.3 (1.3) |
| Completeness (%) | 97.5 (95.9) | 96.0 (76.5) |
| Redundancy | 4.9 (5.0) | 2.8 (2.3) |
| **Refinement** |  |  |
| Resolution (Å) | 34.83-1.95 (2.00-1.95) | 49.45-2.58 (2.65-2.58) |
| No. reflections | 53046 | 87207 |
| *R*_work_ / *R*_free_(%) | 15.6/20.6 | 23.5/27.0 |
| No. atoms | 5101 | 20479 |
| Protein | 4696 | 20130 |
| Ligand/ion | 38 | 256 |
| Water | 339 | 93 |
| *B*-factors | 31.5 | 46.6 |
| Protein | 30.9 | 47.8 |
| Ligand/ion | 24.5 | 37.9 |
| Water | 39.4 | 26.0 |
| R.m.s. deviations |  |  |
| Bond lengths (Å) | 0.005 | 0.005 |
| Bond angles (°) | 1.289 | 1.266 |

*Values in parentheses are for highest-resolution shell.

**Supplementary Table 5. Detailed instrumental parameters of LC-MS.**

| **Parameter setting of the mass spectrometer** | |
| --- | --- |
| **Ion Source:** |  |
| Spray Voltage: 3500 V (pos); 2500 V (neg) | Ion Transfer Tube Temperature: 325 |
| Sheath Gas: 30 Arb | Auxiliary Gas: 10 Arb |
| Vaporize Temperature: 350 | Ion Source Type: H-ESI |
| **Full scan parameters:** |  |
| Scan range: 150-520 *m/z* | Resolution: 60,000 |
| S-lens RF (%): 70 | Automatic Gain Control (AGC) Target: Standard |
| **DIA mode parameters for MS^2^:** |  |
| Precursor Mass Range: | Resolution: 22,500 |
| Isolation Window (*m/z*): 18 | Window Overlap: 1 *m/z* |
| AGC Target: Standard | HCD Collision Energy (%): 15, 30, 60 |
| Loop Control: N | N (Number of Spectra): 7 |
